# Predicting future patterns of land cover from climate projections using machine learning

**DOI:** 10.1101/2024.07.14.603429

**Authors:** Tomasz F. Stepinski

## Abstract

Vegetation plays a crucial role in the Earth’s system, and its characteristics are strongly influenced by climate. Previous studies have investigated the climate-vegetation relationship, often attempting to predict vegetation types based on climate data. Many of them have utilized biome types as proxies for different vegetation forms. Biomes, although widely used, are not always optimal for this task. They are broadly defined, a priori linked to climate, and subject to change over time. This study proposes a novel approach by using the local composition of land cover (LC) categories as descriptors of vegetation types and examines the feasibility of modeling such compositions based on climate data. The investigation focuses on the Sahel region of Africa, which is tessellated into 5 × 5 km square tiles, serving as the basic units of analysis. The independent variable comprises a set of bioclimatic variables assigned to each tile, while the dependent variable consists of shares of each LC category within the tile. The modeling framework involves a set of *n* regressions, one for each LC category. The K-nearest neighbors (KNN) algorithm is employed to ensure that interdependently predicted shares sum up to 100%. The model’s performance is validated using 2010 data, where both climate and LC information are available. The mean absolute value of residuals ranges from 1% to 11%, depending on the LC category. Subsequently, future predictions of LC patterns are made for 2040, 2070, and 2100 using climate projections under IPCC scenarios 370 and 585. A novel visualization technique called synthetic landscape is introduced to visually compare the temporal sequence of predicted LC maps from 2010 to 2100 with similar sequences of biome maps and Köppen-Geiger climate type maps. This comparison highlights overall similarities across all sequences but also reveals some significant differences.

## 1. Introduction

The link between vegetation and climate has been recognized since ancient times. The Greek philosopher Theophrastus (c. 371 – c. 287 BCE) made notable observations about the influence of climate on plant distribution. Categorizing plants based on their forms and structures, he documented variations in vegetation across different climatic regions (Ierodiakonou, 2020). In the early 19th century, Alexander von Humboldt conducted extensive research on the relationship between climate and vegetation during his travels in South America and other regions. His groundbreaking work laid the foundation for understanding the interconnectedness of environmental factors, including climate, soil, and vegetation (Strobl, 2021).

The Köppen-Geiger classification scheme, which is approximately 100 years old but still widely used, is a prime example of the link between climate and vegetation (Köppen, 1936). This classification system parameterizes the boundaries of vegetation zones based on temperature and precipitation (Thornthwaite, 1943). The connection between climate and vegetation is a recurring theme in recent literature, particularly due to the growing interest in climate change and its implications. Readers interested in bibliometric reviews can refer to Afuye et al. (2021) and Li et al. (2021), while Fischlin et al. (2007) provides a comprehensive report on how climate change is expected to impact the future distribution of vegetation zones.

This work constitutes an exploratory study aimed at investigating the feasibility of predicting future land cover patterns from projected climate data. It falls within the realm of data-based modeling of vegetation types and/or vegetation intensity in relation to climate. There exists a substantial body of literature dedicated to such modeling. To provide contextual background for this paper, I will briefly review the existing literature on climate → vegetation data-based modeling.

It is important to note that the term “vegetation” is used here as an overarching concept encompassing various biotic variables. The specific nature of a biotic variable serves as one of several distinguishing factors among different models. Other differentiating factors include the sets of abiotic (climatic and possibly non-climatic) covariates utilized in a model, as well as the algorithm employed in generating the model.

The majority of prior works have focused on models capable of globally mapping different biomes (Levavasseur et al., 2012, 2013; Hengl et al., 2018) and forecasting global shifts between these biome types due to climate change (Bonannella et al., 2023), as well as reconstructing the spatial distribution of biome types in the past (Lindgren et al., 2021). A common thread among these papers is the utilization of the BIOME 6000 dataset (Harrison, 2017) as a training set for inductive modeling. The advantage of the BIOME 6000 dataset lies in its provision of a globally applicable standardized classification. However, a drawback is its relatively small sample size (approximately 8,000 points), which are unevenly distributed across the landmass. Sato and Ise (2022), on the other hand, utilized the 0.5° ISLSCP2 Potential Natural Vegetation Cover grid (Ramankutty et al., 2010) as biomes, thereby circumventing the issue of a small and unevenly distributed training dataset.

In contrast, regional climate → vegetation models make use of locally compiled databases of vegetation types as a training set for inductive modeling. These types are sometimes referred to as biomes (Heubes et al. (2011), Tovar et al. (2013), Tovar et al. (2022)) for the Andes, Zevallos and Lavado-Casimiro (2022) for Peru, or Maksic et al. (2022) for Brazil, and sometimes as vegetation types (Somodi et al. (2017) for Hungary, Hinze et al. (2023) for Europe, Fischer et al. (2019) for Bavaria, Keane et al. (2020) for a region in the US state of Montana, and Liu et al. (2009) for a region in north-east China).

Another branch of literature employs distributions of individual species of vegetation as spatio-temporal biotic variables (Lorena et al., 2008; Franklin et al., 2013; Hanewinkel et al., 2014). Such modeling is based on the concept of the fundamental niche (Peterson, 2001), wherein the fundamental niche of a species is defined as an area with a climate (and other environmental conditions) capable of supporting the species. Algorithmically, this approach differs from the aforementioned methods as each species is modeled separately.

Furthermore, the Normalized Difference Vegetation Index (NDVI) has been utilized as a proxy for a biotic variable (Patil et al., 2017; Bao et al., 2021). Since NDVI is a numerical variable, the model takes the form of regression rather than classification (as seen in the models discussed previously). The model is trained using current NDVI imagery and applied to future climatic conditions to predict future NDVI images.

In principle, modeling vegetation from climate can be achieved through various methods, including process-based mechanistic modeling, expert-based manual modeling, rule-based modeling, and data-based modeling (Hemsing and Bryn, 2012). However, the majority of studies utilize data-based, supervised machine learning (ML) algorithms to induce climate → vegetation models. Different ML algorithms are grounded on different concepts of learning. Notably, the Random Forest (RF) algorithm (Breiman, 2001) has been the most popular choice (Patil et al., 2017; Keane et al., 2020; Lindgren et al., 2021; Zevallos and Lavado-Casimiro, 2022; Hinze et al., 2023) for inducing climate → vegetation models.

Given that these models aim to predict vegetation from climate, the covariates must consist of climatic variables. Most studies commence with 19 bioclimatic variables (O’Donnel and Ignizio, 2012), which are a set of indices highlighting climate conditions most relevant to the physiology of vegetation species. Some studies expand the list of predictors by incorporating other climatic variables, topographic variables, soil variables, etc. For instance, Levavasseur et al. (2012) utilized 43 predictors, Bonannella et al. (2023) employed 72 predictors, and Hengl et al. (2018) utilized 160 predictors. Conversely, some works use only a small subset of bio-climatic variables. For instance, Hinze et al. (2023) utilized only five bioclimatic variables, while Sato and Ise (2022) used average monthly temperatures and precipitation instead of bioclimatic indices.

This paper presents an exploratory study aimed at investigating the possibility of using land cover (LC) data as a “vegetation” variable in modeling the dependence of vegetation on climate. Notably, LC is conspicuously absent from various representations of vegetation used in climate → vegetation models. Land cover categories result from machine learning classification of pixels in remotely sensed multispectral or hyperspectral images (for reviews, see Coppin et al. (2004) and Tewkesbury et al. (2015)). While some land cover categories correspond to natural vegetation, others describe anthropogenic or climate-independent aspects of the land surface. For the purpose of climate → vegetation modeling, the land cover dataset needs to be transformed to represent potential natural vegetation land cover. Unlike biomes, land cover categories are not explicitly defined in terms of climate and represent a lower level of abstraction of vegetation cover. Land cover data are available from multiple sources in the form of global datasets at fine spatial resolution.

Note that there is existing literature on predicting future maps of land cover categories (for reviews, see Liu and Yang (2015), Camacho Olmedo et al. (2018), or Ren et al. (2019)). However, existing land change models are not driven by climate change; instead, they are autoregressive – they extrapolate past temporal trends to predict the future. They are typically used for short-term predictions of deforestation (Bradley et al., 2017) or urban growth (Aburas et al., 2021) where data from several past time stamps are available. Here, my interest lies in non-autoregressive models capable of predicting changes in potential natural land cover over climate change time scales.

LC pixels are too small to reliably predict from climate data alone. Instead, the model presented here predicts the bulk compositions (histograms) of LC categories in a local area defined by a square tile consisting of a number of pixels large enough to be perceived as a pattern, but small enough compared to the total number of pixels in the study area (Cardille et al., 2012; Niesterowicz et al., 2016; Niesterowicz and Stepinski, 2017; Jasiewicz et al., 2018). The predicted quantity is referred to as local land cover composition (LLCC). It is important to note that a full description of a local LC pattern includes the spatial configuration of categories in addition to the LLCC (Wickham and Norton, 1994; Nowosad and Stepinski, 2019). However, predicting pattern configuration is beyond the scope of this investigation.

The goal of this work is to develop a climate → LLCC model capable of forecasting future maps of LLCC under projected climate conditions. A secondary goal is to visualize an actual future land cover (LC) map (not just maps of local abundance of different categories) for direct comparison with the present-day LC map of the same study area. As this is an exploratory study, its aim is to test the hypothesis that predicting future maps of land cover from future projected climate data is possible with reasonable accuracy. Thus, the investigation is limited to the Sahel region in Africa, and the results are not intended as a resource for planning but only to support or refute the hypothesis

## 2. Data

The location of the study area and its geographical context are shown in Fig. 1A. It is situated in Africa between longitudes 18°E and 27°E and latitudes 6°N and 15°N. The study area covers a part of the Sahel region—a transitional zone between the arid Sahara (desert) to the north and more humid regions to the south. Fig. 1B displays a satellite image of the study area (2023 Google Earth, Landsat/Copernicus, image taken on 4/10/2013), while Figs. 1C, D, E, and F show zoomed-in satellite images at locations representative of the four major LC categories present: tree cover, shrubland, grassland, and bare land, respectively.

**Figure 1:**
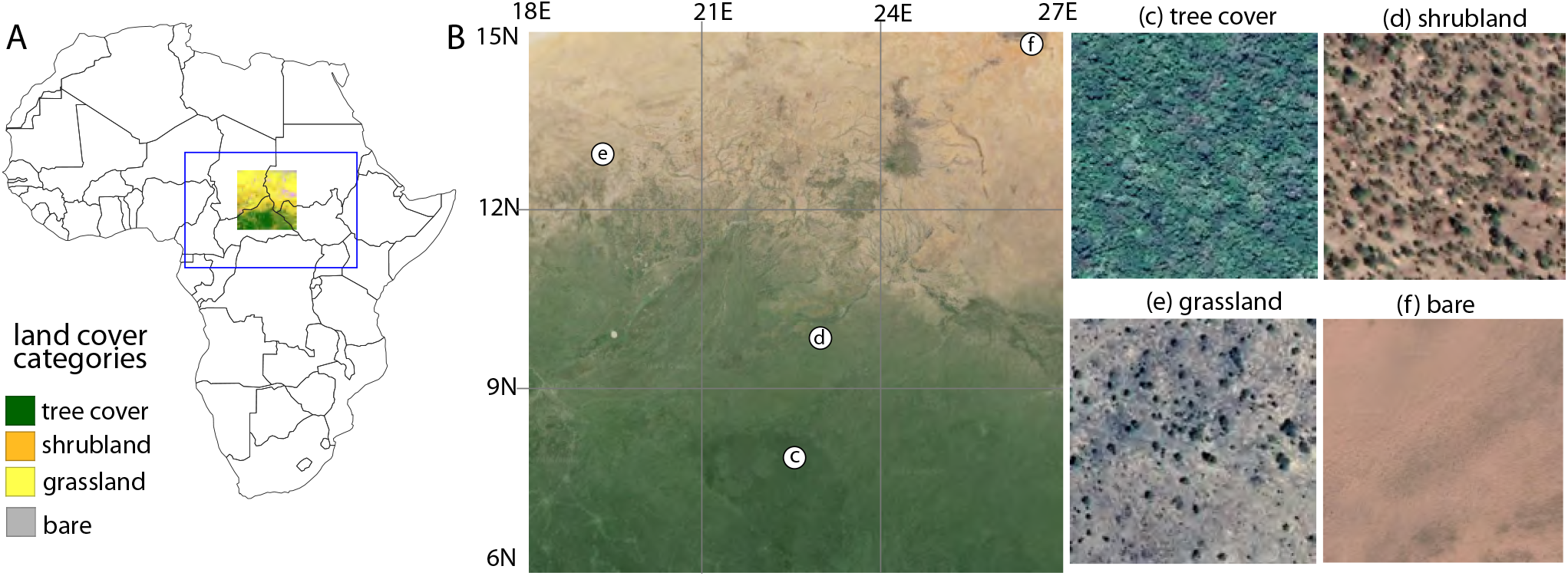
The land cover map depicts the study area. The inset provides a context map showing the location of the study area within the African continent. Grid lines are included to provide locational reference.

The study area is where I will forecast future LLCC. However, future climates may not be restricted to those presently observed within the study area. To partially address this issue, I am using a larger swath of land indicated by a blue rectangle in Fig. 1A (longitudes 10°E – 36°E and latitudes 0°N – 18°N), which encompasses a more diverse sample of present-day climates, as a training area. Present-day climatic and LC data from the training area are used to build a model.

For the LC data, I utilized ESA World Cover 2021 (https://worldcover2021.esa.int), a global product based on Sentinel 1 and 2 data with a resolution of 10 meters per pixel. I resized the LC data to 100 meters per pixel, resulting in a categorical raster covering the extent of the training area with dimensions of 21,600 × 31,200 cells. There are eight different land cover categories present in the study area, of which four are considered “natural vegetation types” (see the legend in Fig. 1A).

For present-day (1981-2010) and future (2011-2040, 2041-2070, and 2071-2100) climates, I utilized data consisting of 19 bioclimatic variables (O’Donnel and Ignizio, 2012) provided by the CHELSA project (Karger et al., 2017). Bioclimatic variables are designed to highlight climate conditions most related to the physiology of vegetation species. For future climates, I considered two IPCC (Intergovernmental Panel on Climate Change) scenarios, 370 and 585. The last two digits of a scenario correspond to additional radiative forcing in 2100 (for example, 85 means 8.5 W/m^2^), and the first digit corresponds to shared socioeconomic pathways: 1 - sustainability, 3 - regional rivalry, and 5 - fossil-fueled development (ref). CHELSA data has a resolution of 1 km, but I resized it to 5 km, resulting in climatic data over the training area as a numerical raster with dimensions of 432 × 624 cells.

## 3. Methods

Methods are divided into several stages: data preprocessing, training the climate → LLCC model, and development of a synthetic land cover map for visualizing the results.

### 3.1. Data Preprocessing

The input to the preprocessing stage consists of two datasets covering the spatial extent of the training set (see the blue rectangle in Fig. 1A). The first dataset is a grid of the LC data, measuring 21,600 × 31,200 cells. Each cell in the LC grid stores a single label from a set of eight possible labels. The second dataset is a grid of climatic data, measuring 432 × 624 cells. Each cell in the climatic grid stores 19 bioclimatic variables.

The LC grid is divided into 50×50 cell tiles. The size of the tile is chosen so that each tile has the same spatial extent as a climatic cell. Thus, a single climate is assigned to a 5×5 km patch of land described by a pattern formed by 2,500 LC cells. The thematic composition of a tile is denoted as *n*_1_, …, *n*_8_, where *n*_*i*_ represents the count of cells in a tile having land cover category *i*. However, over the extent of an area covering the training dataset, there are only four significantly abundant LC categories that are directly predictable from climate: tree cover (*n*_1_), shrubland (*n*_2_), grassland (*n*_3_), and bare land (*n*_4_). These categories are referred to as natural vegetation categories. The remaining categories are grouped into a single category labeled as “other.” After such relabeling, the thematic composition of a tile becomes {*n*_1_, *n*_2_, *n*_3_, *n*_4_, *n*_5_}, where *n*_5_ is the count of cells in the tile that are not directly predictable from climate. The final preprocessing step for the LC data involves relabeling “other” cells in a tile to one of the four natural vegetation labels. Given the small spatial extent of a tile, I simply relabel such cells to the natural vegetation category that is most abundant in the tile. After this transformation, the vegetation part of the dataset can be referred to as potential natural LC. Thus, the LLCC at cell *k* is quantified by 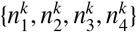.

The nineteen bioclimatic variables are highly correlated with each other. Principal component analysis (PCA) of the bioclimatic variables from the training area reveals that the first four components explain 92% of the variability in the climatic data of the training area. Although these four components could be used to describe the climate, two of them lack a clear interpretation. Moreover, for compatibility, all climatic data, including future projection data, would need to be combined into a single dataset before performing PCA analysis. Instead, I chose to use just four bioclimatic variables: *b*_1_ = bio1 (annual mean temperature), *b*_2_ = bio4 (temperature seasonality), *b*_3_ = bio12 (annual precipitation), and *b*_4_ = bio15 (seasonality of precipitation). Thus, the climate at tile *k* is quantified by four numbers 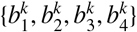

### 3.2. Training the climate → *LLCC model*

A model is trained using a supervised machine learning (ML) technique. Such a model is inductive, as it constructs a prediction function by generalizing from a large number of individual observations. In the present context, observations are in the form of implications 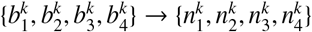, where *k* = 1 to 269568, representing all tiles in the training area. The antecedent (premise) is the local climate, and the consequent (conclusion) is the LLCC.

As the consequent is numerical rather than categorical, an ML model needs to be used in its regression mode. Moreover, the complete model consists of four independently trained regression models, one for each LC category. The major technical challenge is to ensure that the sum of the four predictions is 100% without imposing artificial constraints. Experimentation revealed that this challenge is met by using the KNN algorithm.

The KNN algorithm relies on the idea that tiles with similar climates (in the Euclidean sense) have similar values for the consequent. The free parameter *K* represents the number of tiles that are climatically most similar to a given tile, for which predictions about LLCC are to be made. In this study, I used *K* = 20. For instance, to predict the expected share of grassland pixels 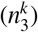 in a tile *k*, the algorithm identifies the 20 climatically most similar tiles from the training set (where both climates and compositions are known from observations) and assigns an average value of their grassland shares as the prediction for 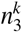. Since the selection of the 20 nearest tiles depends only on the climate, it remains the same for regressing shares of each of the four LC categories. In essence, KNN predicts the shares of a given tile by averaging the shares of 20 training tiles with climates closest to that of the given tile. This guarantees that the sum of all four regressed shares always adds up to 100%.

Machine learning (ML) regression models can be considered as interpolation or extrapolation depending on circumstances. In previous studies related to predicting the future spatial distribution of biomes from climate projections (as discussed in the introduction section), the training set was often small and geographically limited compared to the testing set (the rest of the terrestrial landmass). If this geographical discrepancy extends to the data space, such a training set may not be representative, and an ML model will need to extrapolate, potentially leading to inaccurate predictions (Meyer and Pebesma, 2021, 2022). Spatial crossvalidation was used in previous works to assess the ability of the model to extrapolate, but the results may not be conclusive because cross-validation relies solely on data from the training set, whereas future climates may lie outside that range.

Using LC instead of biomes enables a different approach. I select a training area significantly larger than the intended predicting area, which is located inside the training area (see Fig. 1A). Thus, the range of present climates in the training area is guaranteed to be at least equal, but most likely significantly larger than the range of present climates in the study area. The idea is that even if future climates within a study area extend beyond the envelope of present climates in this area, they may still fall within the envelope of present climates in the training area. This approach minimizes the need for the model to extrapolate from the training data.

Experiments show that it is unnecessary to train the KNN model on data from all 269,568 tiles in the training area. Instead, it is sufficient to choose a random sample of tiles, ensuring they are evenly distributed across the entire training area. Once the sample size reaches 10% of all cells, further increases do not improve the model. The results presented in this paper were obtained using a model induced from a 10% random sample.

### 3.3. Synthetic maps of LC

Given future climate data 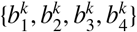 for *k* = 1, 46, 656 tiles within the predictive area of 216 × 216 tiles, a model predicts the future thematic composition of those tiles. For the prediction of a spatial variable to have the most immediate impact, it must be visualized (mapped) in the most expected form. Such a form is a map of LC, not a map of LC composition. It is not possible to construct an actual map of future LC from thematic composition alone because the spatial configuration of cells with different labels within a tile remains undetermined. However, it is possible to construct a synthetic map of LC.

A synthetic map of LC is a patchwork of present-day LC patterns. For a given tile with future composition 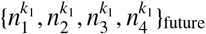, I identify a tile in the present time 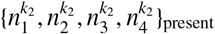 with the most similar composition in the Euclidean sense. I then assign the present pattern at location *k*_2_ as the future pattern at location *k*_1_. The result is a 100 m resolution map of LC with the correct composition at each tile, but likely an incorrect configuration. Such a map lacks continuity at tile boundaries, which, when viewed at full resolution, makes it difficult to discern changing patterns. Downscaling a synthetic map to a resolution of 5 km eliminates the problem while still providing enough details for qualitative visual comparison with similar maps from different times, including a map from the present time.

## 4. Results

This section is divided into two subsections: model validation using the training data and model predictions of future distributions of LLCC compositions and LC maps.

### 4.1. Validation using the training data

The model has been trained as described in Section 3.2. Given the local climate ascribed to tile *k*, encapsulated by four indices 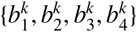, it returns a predicted LC composition of the tile 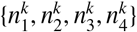. Validation and prediction are performed on 46, 656 tiles located to the study area. All maps in this section depict the entire study area divided into nine sectors by grid lines for locational reference.

The 2010 LC data is used for validating the model. The upper row in Fig. 2A displays four maps depicting the spatial distributions of observed shares of trees, shrubland, grassland, and bare land, respectively. The percentage of tiles occupied by a given LC category is depicted by a color, ranging from deep red (100%) to deep violet (0%). It is evident from the 2010 observations that there is a layered geographical layout of different vegetation types, from trees in the south to grass and bare in the north. The observed bulk composition of the entire study area is as follows: trees - 24%, shrubland - 26%, grassland - 47%, and bare land - 3%.

**Figure 2:**
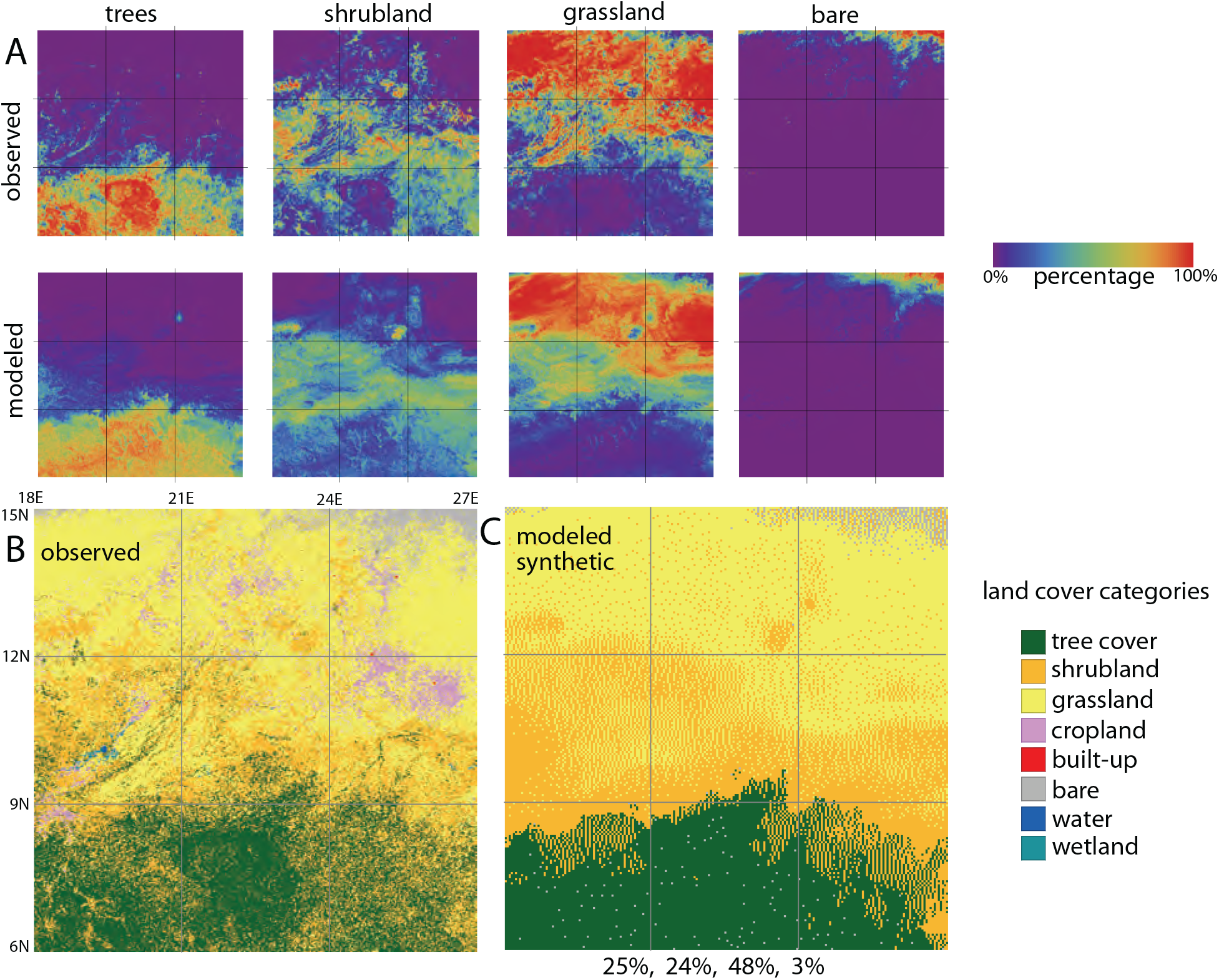
(A) Comparison between LLCD data over the study area as observed in 2010 and as calculated by the model using 2010 climatic data. (B) Map of land cover at 100 m resolution observed in 2010. (C) Synthetic map of land cover at 5 km resolution derived from the composition predicted by the model based on 2010 climatic data. Grid lines provide locational reference. Numbers below panel C represent the bulk composition of the entire study area (tree cover, shrubland, grassland, bare land).

The lower row in Fig. 2A shows LC composition maps based on shares modeled by the 2010 climate. The climate-modeled geographical layout is visually very similar to the observed one. The modeled bulk composition of the entire study area is as follows: trees - 24%, shrubland - 25%, grassland - 48%, and bare land - 3%.

Observed layouts of the composition show sharp boundaries between vegetation types with only small overlaps. In contrast, modeled layouts show somewhat more diffused boundaries. The mean absolute values of the residuals are as follows: trees - 6%, shrubland - 11%, grassland - 10%, and bare land - 1%. This means that, for example, on average (over all tiles in the study area), the discrepancy between observed and modeled shares of trees is 6%.

Fig. 2B shows a comparison between an observed and predicted synthetic maps of LC in 2010. The observed LC map has a resolution of 100 m and displays eight different LC categories, of which four categories (cropland, built-up, water, and wetland) are not predictable from climate. The predicted LC map is synthesized as described in Section 3.3 and has a resolution of 5 km. It displays only four LC categories that are predictable from climate. Synthetic maps serve as visualizations; quantitative analysis should be performed using predicted compositions (depicted in Fig. 2A).

### 4.2. Future LLCC predictions

Fig. 3 shows maps depicting predicted spatial distributions of shares of trees, shrubland, grassland, and bare land in 2040, 2070, and 2100 under two different IPCC scenarios. As in Fig. 2A, color indicates the percentage of a tile’s area occupied by a given LC category, ranging from deep red (100%) to deep violet color (0%).

**Figure 3:**
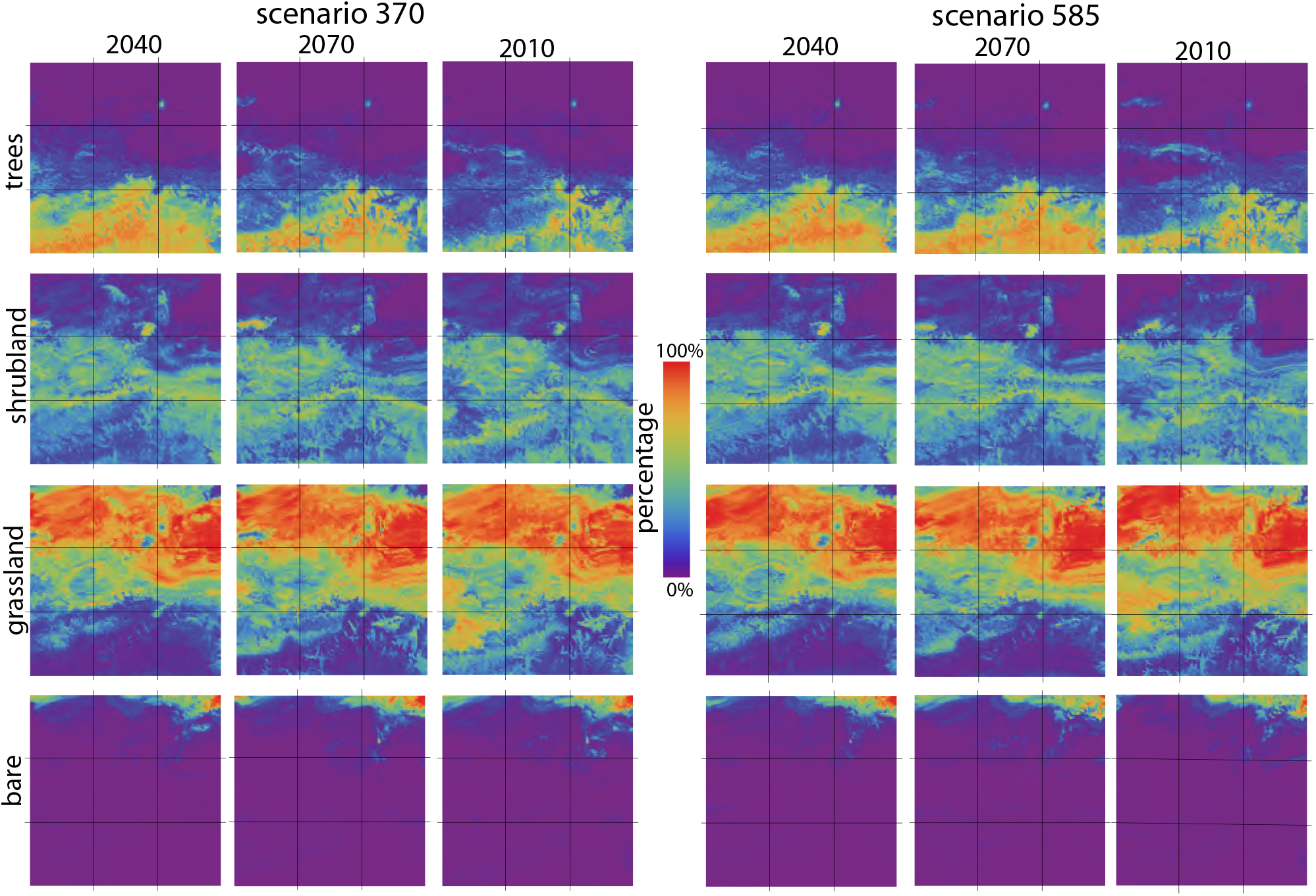
Future LLCC predictions for 2040, 2070, and 2100. (Left panels) Predictions using climate projections from the IPCC 370 scenario. (Right panels) Predictions using climate projections from the IPCC 585 scenario. Grid lines provide locational reference.

The results shown in the first three left columns in Fig. 3 correspond to scenario 370. As explained in the introduction, the first digit, 3, indicates a scenario or shared socioeconomic pathway (SSP). SSP3 assumes a resurgent nationalism, concerns about competitiveness that push countries to increasingly focus on domestic issues and giving environmental concerns a low priority. The additional radiative forcing achieved by 2100 is 7 W/m^2^. Each column shows the predicted composition in a different year, while each row shows the temporal evolution of predicted shares for a given LC category. Prepending each evolutionary series with a corresponding 2010 frame from the lower row in Fig. 2A illustrates 90 years of landscape evolution due to projected climate change.

The most striking result of the prediction is the dying out of the southern zone of tree cover, which becomes pronounced after 2040. The shrubland category, which in 2010 was present south of 9°N only in the westernmost sector, progressively increases its share over time in all southernmost sectors at the expense of trees. The grassland category, virtually absent south of 9°N in 2010, expands southward starting in 2040, especially in the eastern sectors. The area covered by the bare land category does not show significant spatial change from 2010 to 2100.

The results shown in columns four to six in Fig. 3 correspond to scenario 585, also referred to as SSP5. SSP5 assumes increasing reliance on competitive markets and innovation to produce rapid technological progress. As a result, the exploitation of fossil fuel resources and the adoption of energy-intensive lifestyles around the world increase. The additional radiative forcing achieved by 2100 is 8.5 W/m^2^. The evolution of the spatial distribution of trees, shrubland, grassland, and bare land due to climate change under scenario 585 is not markedly different from their evolution under scenario 370. The disappearance of trees in the southernmost parts of the study area predicted by the model under scenario 585 is slower than predicted by the model under scenario 370. Fig. 4 displays synthetic maps of land cover constructed from predictions of LLCC, offering a simplified yet direct visualization of the overall evolution of vegetation in the study area under two climate change scenarios. The retreat of the tree zone is clearly evident in both scenarios, although it occurs faster under scenario 370. Shrubland fills the space formerly occupied by trees, and there is a noticeable southern progression of grassland in the eastern sectors of the study area. The changes in the bulk composition of land cover categories across the entire study area quantify the net loss of tree cover from as much as 24% in 2010 to as little as 14% in 2100, alongside the gain of grassland cover from as little as 38% in 2040 to as much as 57% in 2100.

**Figure 4:**
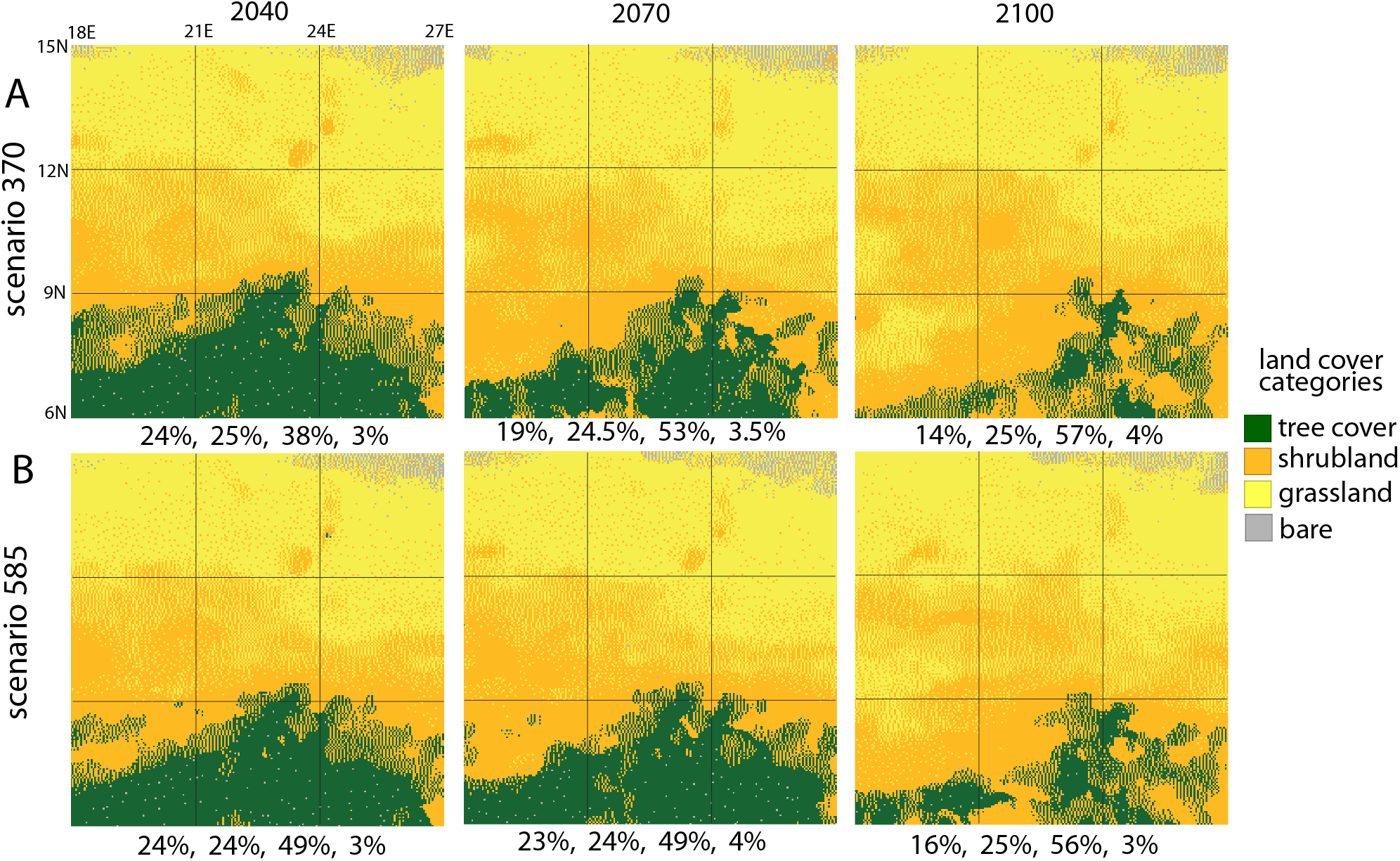
Synthetic maps of land cover in 2040, 2070, and 2100, based on predictions of LLCC under scenario 370 (upper row) and 585 (lower row). The numbers below each map represent the bulk composition for the entire study area: trees, shrubland, grassland, bare land.

## 5. Disussion

In this section, I compare my results to previously made predictions of vegetation redistribution due to projected climate change in the location of my study area and discuss the impact of the employed technical solutions on the plausibility of the prediction.

For comparison, I selected predictions made by Bonannella et al. (2023) and Beck et al. (2023). Bonannella et al. predict the future distribution of biomes for the periods 2040-2060 and 2062-2080 under three climate change scenarios (Representative Concentration Pathways (RCP) 2.6, 4.5, and 8.5). They publish their projections for the entire terrestrial landmass, using the inductive ML algorithm (Random Forest) and training data from 1979-2013.

Beck et al. predict global future Köppen-Geiger maps using different climate projection scenarios. While these are maps of climate types, the Köppen-Geiger classification is based on expressing the boundaries of vegetation zones in climatic terms. Notably, in the location of the study area, Köppen-Geiger climate types have names associated directly with vegetation, such as tropical savanna, arid desert, or arid steppe, suggesting they can be considered vegetation zones (see Table 4 in Denk et al. (2013)).

Fig. 5 compares the temporal changes of three different proxies of vegetation due to climate forcing. In my method, the calculated variable is LLCC. However, for comparison with other methods, consecutive snapshots of synthetic LC maps are shown in row A of Fig. 5. Consecutive snapshots of biome maps (Bonannella et al.) are shown in Fig. 5B. Finally, consecutive snapshots of Köppen-Geiger maps (Beck et al.) are shown in Fig 5C. The climate change scenario for sequences A and C is 370, and it is 4.5 for sequence B. Note that the categories displayed by different methods are not directly comparable (see corresponding legends in Fig. 5). All three temporal sequences exhibit a layered (south-to-north) structure of vegetation zones consistent with the climatic changes in the region. However, the meanings and extents of the layers differ. In sequences A and B, the southernmost layer is characterized by forests, while in sequence C, it is identified as a tropical savanna. According to Denk et al. (2013), the vegetation in the KG climate type “tropical savanna” consists of semi-evergreen and deciduous forests. Moreover, the spatial extent of the southernmost layer varies from minimal in sequence B to very extensive in sequence C. Moving northward, the northernmost layer is occupied by grassland (and some bare land) in sequence A, a steppe biome (covered by grassland) or xerophytic woods/scrubland biome in sequence B, and an arid desert KG type (semi-desert and desert vegetation) in sequence C. The middle layer is covered by shrubland in sequence A, a tropical savanna biome (tall grasses and occasional trees) in sequence B, and an arid steppe KG type (dry woodlands or grassland) in sequence C.

**Figure 5:**
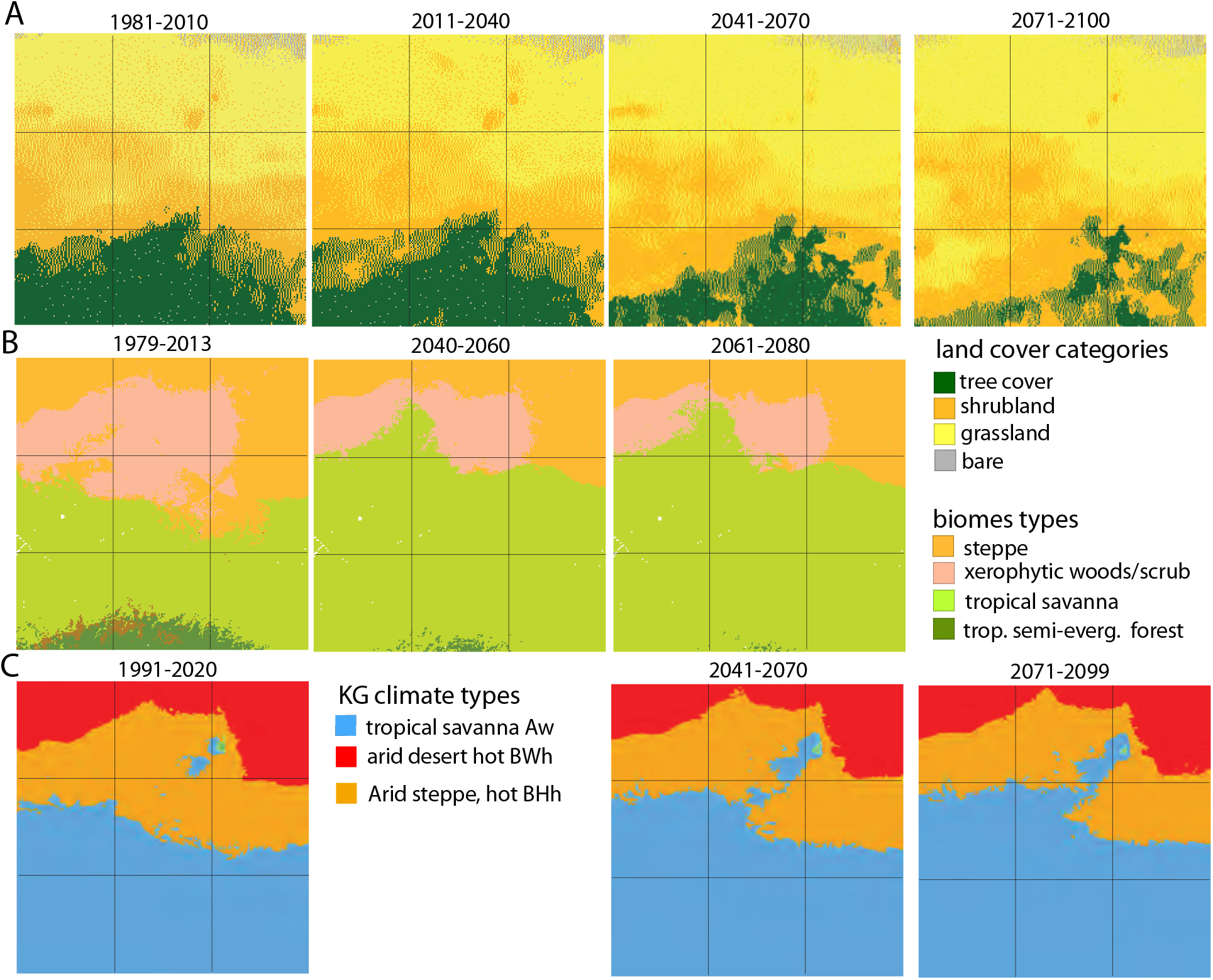
Maps of landscape pattern types (LPTs) in the testing region. The upper right section shows the ground truth map of LPTs and the legend. The remaining three sections pertain to predictions made by three classification algorithms. From the upper-left map clockwise: predicted LPT map, predicted potential natural vegetation map, ground truth map with unpredictable tiles eliminated, predicted map with unpredictable tiles eliminated. See the main text for details.

The most significant differences among the three sequences lie in the representation of tree cover in the southernmost part of the study area. Sequence A depicts a notable presence of tree cover, while sequence B shows a much smaller extent of tree cover in this region. In contrast, sequence C indicates the absence of tree cover entirely within the study area. Additionally, there are evolutionary discrepancies among the sequences. In sequence A, all three vegetation zones (excluding bare land) are migrating south, although at different rates. The retreat of tree cover, occurring at the fastest pace, is followed by the southward movement of shrubland, and grassland, predominantly in the western part of the study area. In sequence B, while the forest retreats southward, the tropical savanna exhibits a slight northward progression. Similarly, in sequence C, the tropical savanna also experiences a slight northward advancement.

The observed discrepancies between LLCC maps, biomes, and KG classifications may partly stem from the specificity of LC definitions compared to the broader definitions of biomes and KG types. However, this disparity cannot solely account for differences in forest coverage. While synthetic LC maps may overstate the presence of forests, a thorough examination of Fig. 2A and Fig. 3 unequivocally demonstrates the current existence of forests in the southernmost region of the study area, with predictions indicating a diminishing extent in the future. Satellite imagery (Fig. 1C) and the land cover map (Fig. 2B) offer definitive evidence of the forest’s present existence in this region.

None of the sequences presented support the hypothesis of a future “greening of the Sahara” (for a comprehensive review, refer to Pausata et al. (2020)). This hypothesis hinges on climate models predicting increased precipitation in the region during June, July, and August, particularly under scenario 585. However, it overlooks simultaneous temperature increases. The minor northward shifts observed in vegetation zones in sequences B and C do not constitute a “greening” phenomenon.

As outlined in the introduction, this study aimed to assess the feasibility of predicting future LLCC from climate projections. To streamline calculations, several simplifications were made, including the following key assumptions:

- Representativeness of the training set for future climates: It was assumed that the training set adequately represents future climates without prior verification of this assumption.
- Representation of climate variables: Climate was represented using only four variables.
- Regression model selection: The regression model was based on the KNN algorithm to ensure result consistency.
- Restriction of the study area: The study area was confined to specific climatic conditions.

Meyer and Pebesma (2021) argued that it’s crucial to establish an “area of applicability” (AOA) for the model given a training set. In this context, the AOA comprises locations where the minimum distance in climate space between the climate forcing a prediction and the climates in the training set is smaller than a threshold. Put simply, predictions of vegetation variables should only be made if the climate driving the prediction is sufficiently similar to some climates present in the training dataset. Adhering to this principle minimizes instances where the model needs to extrapolate, thereby enhancing the reliability of predictions.

I conducted a visual check to assess adherence to the AOA rule. Fig. 6 displays visual AOA tests for all models presented in this study. To facilitate visualization, the climatic space is reduced to two dimensions: total annual precipitation and annual mean temperature.

**Figure 6:**
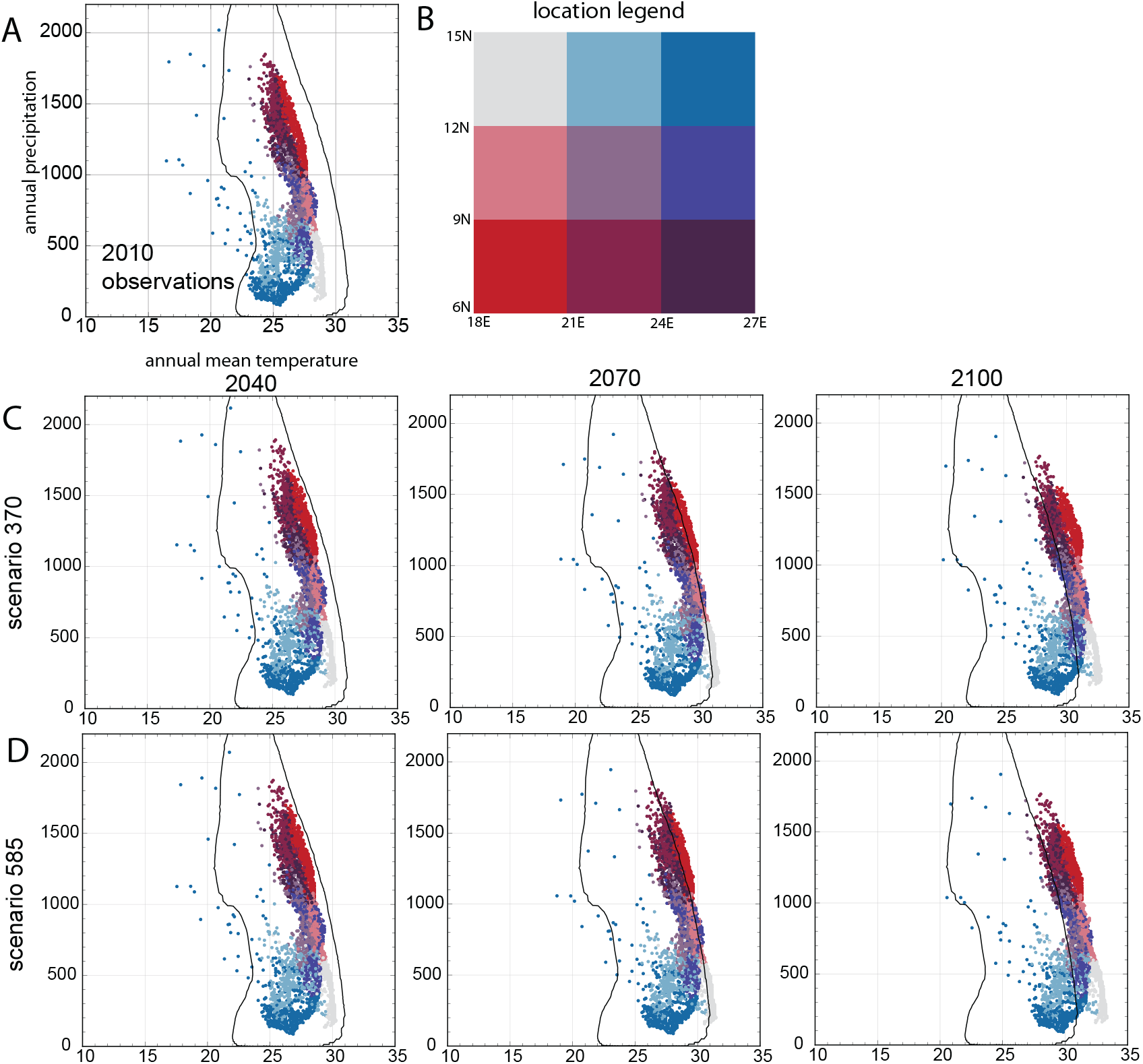
Visual a posteriori tests of adherence to the area of applicability (AOA) for all predictions. The inside of a black contour is an AOA. Points represent values of mean annual temperature and total annual precipitation on a 10% random, uniformly distributed sample of tiles. Points’ colors indicate geographical location (B). (A) Test for observed data in 2010. (C) Tests for future predictions under climate scenario 370. (D) Tests for future predictions under climate scenario 585.

The black contour on each temperature-precipitation diagram represents the envelope of all (temperature, precipitation) values in the training set. Colored points denote a 10% random, evenly distributed sample of (temperature, precipitation) values from among the 46,656 tiles in the study area. Each point’s color indicates its spatial location, categorized into quadrants (see Fig. 5B). Ideally, all points should fall within the envelope.

This criterion is met for the 2010 observations and models for 2040. Some outliers to the left of the envelope correspond to tiles in the Marrah Mountains—a range of volcanic peaks with elevations of up to 3,042 meters above sea level, where temperatures are significantly lower than in the rest of the study area.

In 2070 and especially in 2100, climate projections for many tiles in the study area extend beyond the envelope on the right side. This indicates that predicted future temperatures in these tiles exceed those in any tiles in the training area in 2010. In fact, the climatic conditions for some of these tiles are not currently observed anywhere on Earth. Consequently, predictions of LLCC values for 2100 and even 2070 require extrapolation. It’s noteworthy that the out-of-envelope conditions are primarily caused by rising temperatures, as changes in precipitation values occur at a much smaller relative rate. The most affected quadrants are those along the western edge of the study area, which explains why the southern edge of grassland migrates farther south, particularly in the western part of the study area (as observed in Fig. 2).

Formally, predictions made outside of the AOA should be flagged as uncertain. However, informally, in this case, the prediction is likely to be correct. Extrapolating predictions based on correct precipitation and lower temperatures still results in dominant grassland. Considering the available LC categories for higher temperatures, bare land is the only other viable option. However, given the values of precipitation, it is not a likely outcome. It’s important to note that this discussion is solely based on the mean annual temperature and annual precipitation, disregarding the seasonality of temperature and precipitation—the other two climate variables used in my model.

Does using only four bioclimatic variables (bio1, bio4, bio12, bio15) skew the results? To address this, I conducted sample calculations using all 19 bioclimatic variables, and the results were not significantly different. However, this finding may not generalize to other study areas. One argument against using all 19 bioclimatic variables is their high correlation. Additionally, because the regression is performed using the KNN algorithm, which calculates distances in climatic space, assigning weights to 19 variables would be necessary, as they are not all equally important (Meyer and Pebesma, 2021). Here, I opted to use the same weight for all four bioclimatic variables.

The usage of the KNN algorithm was chosen because it inherently ensures that all predicted shares in each tile sum up to 100%. While other algorithms, such as the Random Forest algorithm, may yield slightly better accuracy for models of individual LC categories, they lead to a departure from the sum of predictions totaling 100%. Addressing the technical challenge of using multiple regression models under constraints, beyond just KNN, could be a promising avenue for future research.

Moreover, it’s important to acknowledge that the selected study area is climatically straightforward. Therefore, it is crucial to further test the method in diverse locations. LC data of the same origin and resolution as used in this paper are available for the entire terrestrial landmass. Additionally, exploring the possibility of utilizing LC data with higher thematic resolution, such as the ESA CCI LC global dataset (available at https://maps.elie.ucl.ac.be/CCI/viewer/), could provide valuable insights for such investigations.

## 6. Conclusions

I have demonstrated that predicting future land cover patterns from climate change projections is feasible. As a proxy for vegetation, LC brings several new possibilities compared to biomes in the realm of future prediction.

Consider the relationship between LC categories and biomes. LC categories define the physical characteristics of the Earth’s surface, encompassing both natural elements (such as forests, grasslands, or shrublands) and anthropogenic features (such as urban areas or agricultural lands). On the other hand, biomes classify largescale ecological communities based on climate, vegetation, and other environmental factors. Thus, the relationship between biomes and LC categories can be likened to that between molecules and atoms in chemistry. Just as molecules are formed by combining different types of atoms in various configurations, biomes combine different LC categories in diverse spatial arrangements. For example, a “tropical savanna” biome may consist of various LC categories such as trees, shrubs, and grass.

From this relationship, it follows that while biomes may serve as suitable areal units for mapping changes in the spatial distribution of vegetation zones on a global scale, LC categories provide more detailed predictions that may be necessary on a regional scale. Additionally, LC categories, being “atomic” units, remain timeless; they are the same now as they will be in the future. In contrast, biomes are defined for the present time, and as the climate changes, some present biomes may become extinct while new ones may appear. This situation is not ideal from the perspective of machine learning inductive modeling, which can only predict future biomes from the set of present biomes.

Finally, future biome-based predictions manifest as categorical maps with hard boundaries between zones (see Fig. 5B). In contrast, the method presented in this paper predicts compositions of LC categories over small areal units (tiles), thus providing quantitative predictions. As a result, synthetic maps of LC (see Fig. 5A) depict not only zones dominated by a single LC category (at the scale of a tile) but also transitional zones between them. Although I utilized 5 × 5 km tiles, the LC data (with a resolution of 10 m) can support quantitative predictions on the scale of 1 km^2^ – the current limit on spatial resolution of climate projections.

To the best of my knowledge, this is the first paper addressing the feasibility of predicting future LC patterns from climate projections. As I pointed out in the discussion section, individual elements of the method may require further development to enhance its efficacy.

## Supporting information

supplement data and code to reproduce results

