## supplement data and code to reproduce results for "Predicting future patterns of land cover from climate projections using machine learning": readme.rtf

This supplement contains a code and all the data needed to reproduce the predictions described in the paper "Predicting future patterns of natural land cover from climate projections using machine learning."
Content:
trainingAndPredicting.ows     --  code to train the model and to use it to make predictions.
This file is a Orange Data Mining workflow. it was was written using version 3.29.3 of Orange. Orange is a visual grad-and-drop interface to Python machine learning codes. It is an open-source (https://orangedatamining.com/)  thus allows anyone, including those without coding skils, to reproduce results presented in the paper. 
trainingSet.csv  - input file containing training data. It has 269,568 rows and 8 columns.  Rows are the tiles in the training set. The first tile is in the upper left corner, other files continue row-wise, the last tile is in the bottom right corner.  Data in a tile is {b1,b2,b3,b4,n1,n2,n3,n4}. bioclimatic data is in original units, shares are in percentages, sum of shares in a tile adds to 100%.
2010ClimatesAndLLCC.csv   --  input file containing observed climatic and land cover data in 2010 over the study area. It has 46,656 rows and 8 columns. Rows are the tiles in the study area. Data is organized in the same way as in trainingAndPredicting.csv.
climate2040_370.csv
climate2040_585.csv
climate2070_370.csv
climate2070_585.csv
climate2100_370.csv
climate2100_585.csv
These six files are climatic input consisting of projections to years 2040, 2070, and 2100 under two different scenarios. Each data has 46,656 rows and 4 columns.  Rows are the tiles in the study area.  Data is a tile is {b1,b2,b3,b4}.
predictedComposition2010.csv  -- output data obtained by applying the model to observed climates in 2010.  It has 46,656 rows and 4 columns. Data for each tile is {n1,n2,3,n4}, predicted values of shares for tree cover, shrubland, grassland, and bare land. Comparison of this prediction with observed shares is used for accuracy assessment.
PredictedComposition2040_370.csv
PredictedComposition2040_585.csv
PredictedComposition2070_370.csv
PredictedComposition2070_585.csv
PredictedComposition2100_370.csv
PredictedComposition2100_585.csv
These six files are outputs consisting of predicted values of composition in each tile (tree cover, shrubland, grassland, and bare land) in years 2040, 2070, and 2100 under two different scenarios. Each data has 46,656 rows and 4 columns.  Rows are the tiles in the study area.  Data is a tile is {n1,n2,n3,n4}.
